# The endosomal protein sorting nexin 4 is a novel synaptic protein

**DOI:** 10.1101/860940

**Authors:** Sonia Vazquez-Sanchez, Miguel A. Gonzalez-Lozano, Alexarae Walfenzao, Ka Wan Li, Jan R.T. van Weering

## Abstract

Sorting nexin 4 (SNX4) is an evolutionary conserved protein that mediates recycling from the endosomes back to the plasma membrane in yeast and mammalian cells. SNX4 is expressed in the brain, but its neuronal localization and function have not been addressed. Using a new antibody, endogenous neuronal SNX4 co-localized with both early and recycling endosomes, similar to the reported localization of SNX4 in non-neuronal cells. Neuronal SNX4 was accumulated in synapses, and immuno-electron microscopy revealed that SNX4 was predominantly localized to presynaptic terminals. Acute depletion of neuronal SNX4 using independent shRNAs did not affect the levels of the canonical SNX4-cargo transferrin receptor. Explorative mass spectrometry showed that each SNX4-targetted shRNA resulted in a reproducible and distinct proteome and that synaptic communication-related proteins were downregulated upon expression of the three shRNAs. The identification of SNX4 as a novel synaptic protein indicates a selective demand for SNX4 dependent sorting in synapses.

## INTRODUCTION

Sorting nexin 4 (SNX4) is an evolutionary conserved protein that mediates endosomal recycling from endosomes back to the plasma membrane^1,2^. SNX4 is a member of the sorting nexin family (SNX) characterized by a phosphatidylinositol 3-phosphate binding domain (phox homology (PX) domain)^3^, which is necessary for peripheral membrane localization^4,5^. More specifically, SNX4 is part of the SNX-BAR subfamily characterized by a carboxy-terminal Bin/|Amphiphysin/Rvs (BAR) domain, which binds to curved membranes upon dimerization^4,6^. SNX4 forms tubules that emanate from the endosomes during the RAB5-Rab7 transition (early endosome to late endosome) and during Rab4-RAB11 transition (early recycling endosome to endosome recycling compartment)^7^. Hettema et al. (2003) showed that silencing the yeast homologue of SNX4 (Snx4p) decreases Scn1p (an exocytic v-SNARE) in the plasma membrane and increases Scn1p degradation at the vacuole (the lysosome equivalent in yeast)^8^. In HeLa cells, a similar SNX4 pathway has been observed: SNX4 recycles back to the plasma membrane the transferrin receptor (TfnR), an iron-transporting receptor located to the plasma membrane, avoiding lysosomal degradation^9^.

SNX4 is expressed in the brain, and SNX4 protein levels are 70% decreased in the brains of severe Alzheimer’s disease (AD) cases^10^. Beta-secretase 1 (BACE-1) is an enzyme involved in proteolytic processing of the amyloid precursor protein (APP), which leads to the formation of the pathological aβ peptide in AD^11^. Two recent studies indicate that SNX4 dysregulation mis-sorts BACE-1 to late endosomal compartments, which impacts aβ production^10,12^. Hence, neuronal SNX4 dysregulation might be a process involved in aβ production and AD etiology.

While SNX4 function is associated with pathological mechanisms in the brain, the physiological role and subcellular distribution of SNX4 in neurons is currently unknown. Here, we characterized the localization of endogenous SNX4 in primary mouse neurons using a new antibody. Endogenous SNX4 partially co-localized with both early and recycling endosome markers, which is in accordance with the previously established role of SNX4 in non-neuronal cells. In contrast, its depletion did not decrease the levels of the best described SNX4 cargo, the transferrin receptor^9^. Neuronal SNX4 accumulated specifically in synaptic areas with a predominant localization to presynaptic terminals, suggesting that SNX4 fulfills a specific role in this compartment. Using three different shRNAs, explorative mass spectrometry revealed that synaptic communication-related proteins were downregulated upon SNX4 depletion. However, each SNX4-targetted shRNA resulted in a reproducible and distinct proteome, which suggests potential off-target effects.

## RESULTS

### Novel SNX4 antibody specifically labels endogenous mouse SNX4 on western blot and immunocytochemistry

Commercially available antibodies against SNX4 only detected mouse SNX4 by western blot. In order to characterize the subcellular localization of endogenous SNX4 in mouse neurons, we have developed a novel antibody. This novel antibody was designed against the N-terminal region of mouse SNX4 in collaboration with Synaptic Systems (Cat. No. 392 003) (Supplementary Figure S1). To confirm that the novel antibody specifically detects SNX4, we developed three independent shRNAs against SNX4, and rescue constructs. Cortical mouse neurons were lentiviral infected at DIV3 with the rescue SNX4 constructs (R1, R2 and R3), and at DIV7 with the three shRNA against SNX4 (shSNX4-1, shSNX4-2, and shSNX4-3) and the shRNA control (Control). At DIV14-15 the neurons were lysed or fixed, and SNX4 levels were evaluated using western blot and immunocytochemistry (Figure 1). Using western blot, the antibody detected a protein of ~50 kDa, which corresponds with the size of SNX4. This band was decreased when using the three independent shRNAs against SNX4 and these levels were restored when the shRNA against SNX4 was combined with the SNX4 rescue constructs (Figure 1a-c). The same ~50 kDa band was observed using two commercially available antibodies (Supplementary Figure S1). Our new antibody also detected a lower band of ~30 kDa in all samples that remained unaffected in the neurons expressing shRNA against SNX4, suggesting that this antibody mediates unspecific binding of protein (Figure 1b). Homogenates of different brain regions were studied with this antibody to gain resolution on SNX4 distribution in the brain. The ~50 kDa band appeared in all homogenates tested (Supplementary Figure S2), suggesting that SNX4 is ubiquitously expressed in the mouse brain.

**Figure 1:**
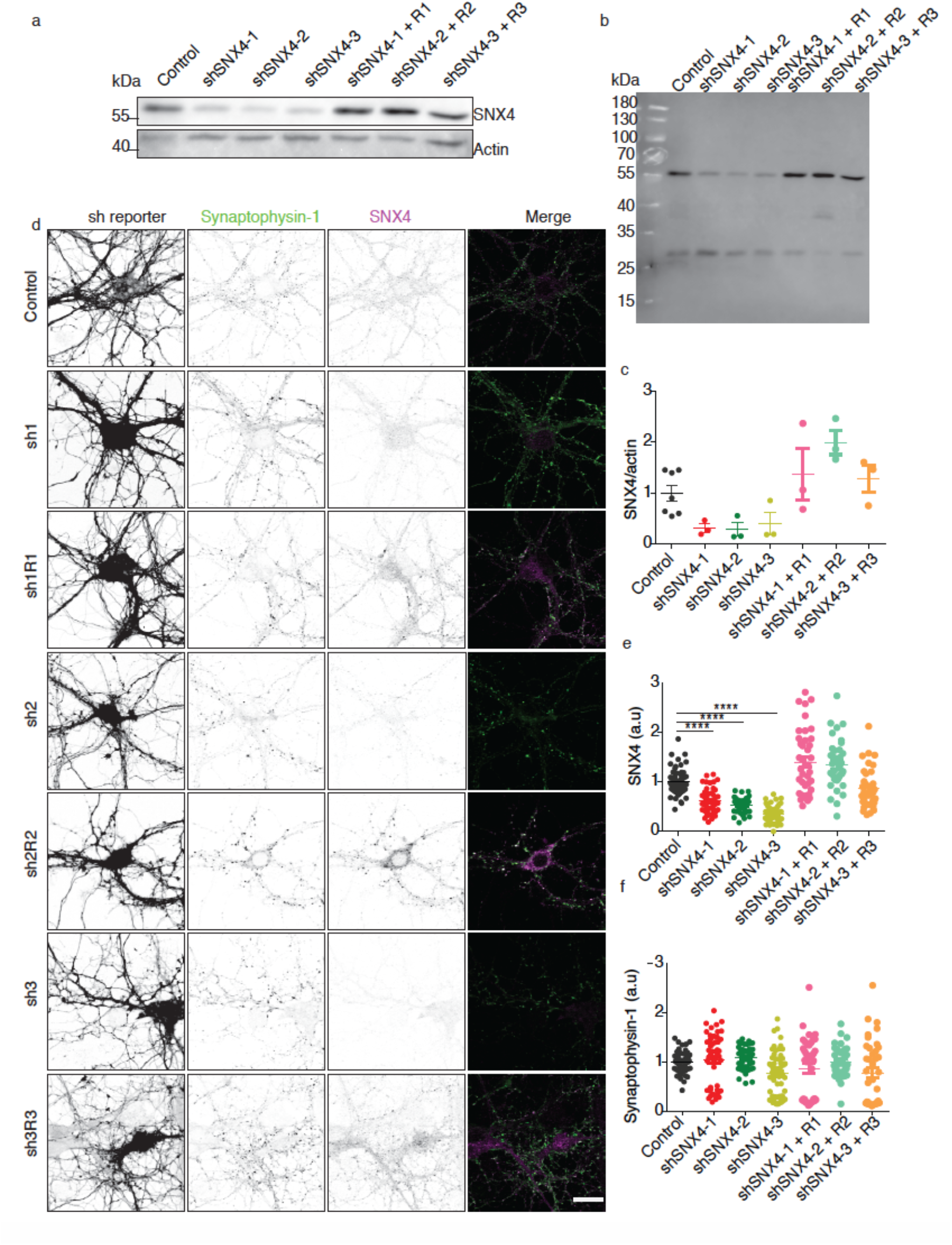
Novel SNX4 antibody specifically labels endogenous mouse SNX4 by western blot and immunocytochemistry. **(a)** Representative SNX4 and actin western blot of control neurons, neurons transfected with shRNAs against SNX4 and neurons transfected with the shRNA and its rescue construct (description of the constructs in materials and methods). Original uncropped actin blot is shown in Supplementary Figure S3. **(b)** Original uncropped blot for SNX4 shown in (a). **(c)** Quantification of SNX4 levels normalized to actin in western blot. Values are presented as a ratio compared to the control condition. (N=3 blots/animals). **(d)** Confocal microscopy images of neurons infected with control shRNA, the three shRNAs against SNX4 and its respective rescue constructs. Left, mCherry signal reporting the transfection of the shRNAs coding sequences. Middle, synaptophysin-1 labelling. Right, SNX4 labelling. (n=50±13 fields of view, N=4±1 animals). Scale bar=20 μm. **(e)** Quantification of synaptophysin-1 staining intensity relative to control. **(f)** Quantification of SNX4 staining intensity relative to control. Detailed information (average, SEM, n and statistics) is shown in Supplementary Table S1.

To test the application of the new antibody in immunocytochemistry, DIV15 cortical neurons were stained with SNX4 and sypnaptophysin-1 antibodies (synaptic marker used as control) (Figure 1d). Expressed mCherry was used as a transfection reporter of all shRNA constructs (both shRNA control and against SNX4). The SNX4 antibody signal showed a punctate pattern in murine primary neuron cultures (Figure 1d). The total SNX4 signal intensity was decreased upon SNX4 knockdown and restored when the shRNA was combined with the rescue constructs, while the total synaptophsyin-1 intensity remained unchanged upon modulation of SNX4 levels (Figure 1d-f). This indicates that the antibody signal was specific for SNX4 in immunocytochemistry. Together, these data confirm that SNX4 is detected using the novel antibody both in western blot and immunocytochemistry.

### SNX4 is located at neuronal early and recycling endosomes

SNX4 has been found colocalizing with early and recycling endosomal markers in HeLa cells, where it coordinates recycling from early endosomes to the plasma membrane through the recycling endosomes^9^. To test if SNX4 also colocalizes with these endosomal makers in neurons, hippocampal mouse neurons at DIV14-15 were fixed and labelled for endogenous SNX4, RAB5 (early endosome marker) and RAB11 (recycling endosome marker). The specificity of the SNX4 localization to these endosome markers was assessed by including a SNX4 knockdown (shSNX4-2) control. Upon SNX4 depletion, the total neuronal levels of RAB5 were 19% decreased (Figure 2a, b). SNX4 colocalized with RAB5 signal (Pearson’s coefficient 0.58) and dropped significantly upon SNX4 depletion (Pearson’s coefficient 0.41, Figure 2 a, c). Expression of shSNX4-2 decreased the levels of SNX4, but it did not affect the levels of RAB11 (Figure 2d, e). The Pearson’s coefficient for the colocalization between RAB11 signal and SNX4 signal was 0.45 which dropped to 0.31 upon SNX4 depletion (Figure 2 d, f). These data indicate that endogenous SNX4 is located to early and recycling endosomes in primary mouse neurons.

**Figure 2:**
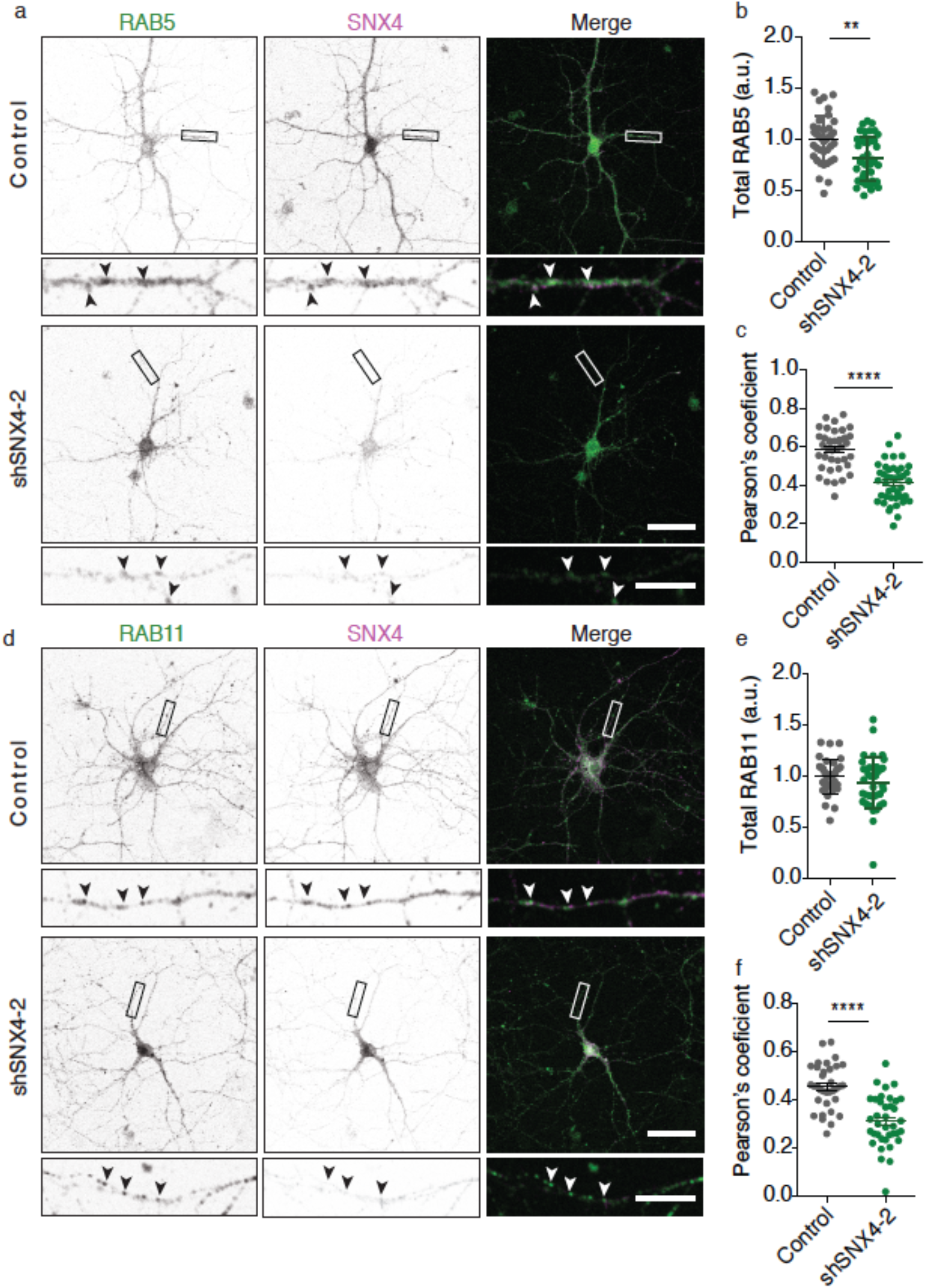
Neuronal SNX4 is located to early and recycling endosomes. **(a)** Confocal microscopy images of control and SNX4 knockdown neurons labelled for RAB5 and SNX4. Merge image of RAB5 (green) and SNX4 (magenta). (n=21±2 neurons, N=3 animals). Scale bar of the neuron image = 50 μm, scale bar of the zoomed neurite=5 μm. **(b)** Quantification of total RAB5 levels in the neuron normalized to control. **(c)** Pearson’s coefficients for the co-localization of RAB5 and SNX4 in neurites. **(d)** Confocal microscopy images of control and SNX4 knock down neurons immunolabelled with RAB11 and SNX4. Merge image of RAB11 (green) and SNX4 (magenta). (n=38±1neurons, N=3 animals). Scale bar of the neuron image=50 μm, scale bar of the zoomed neurite=5 μm. **(e)** Quantification of total RAB11 levels in the neuron normalized to control. **(f)** Pearson’s coefficients for the co-localization of RAB11 and SNX4 in neurites. Detailed information (average, SEM, n and statistics) is shown in Supplementary Table S1.

### SNX4 is located to synaptic terminals

The SNX4 signal appeared to be localized at synapses (Figure 1). To confirm this, we analyzed colocalization between SNX4 and synaptic markers. Control and SNX4 knockdown hippocampal neurons at DIV14-15 were fixed and labelled for SNX4 and synaptophysin-1 (synaptic marker) (Figure 3a). The Pearson’s coefficient for the colocalization between synaptophysin-1 signal and SNX4 signal was 0.71 but this value dropped to 0.55 upon SNX4 depletion (Figure 3b). Further analysis of Figure 1d data showed that SNX4 signal in synaptophysin-1 puncta was decreased upon SNX4 knockdown and restored when the shRNA was combined with the rescue constructs (Supplementary Table S1). This colocalization between synaptic markers and SNX4 was confirmed using an independent antibody against VGLUT1 (Supplementary Table S1). These data demonstrate SNX4 is targeted to synaptic locations.

**Figure 3:**
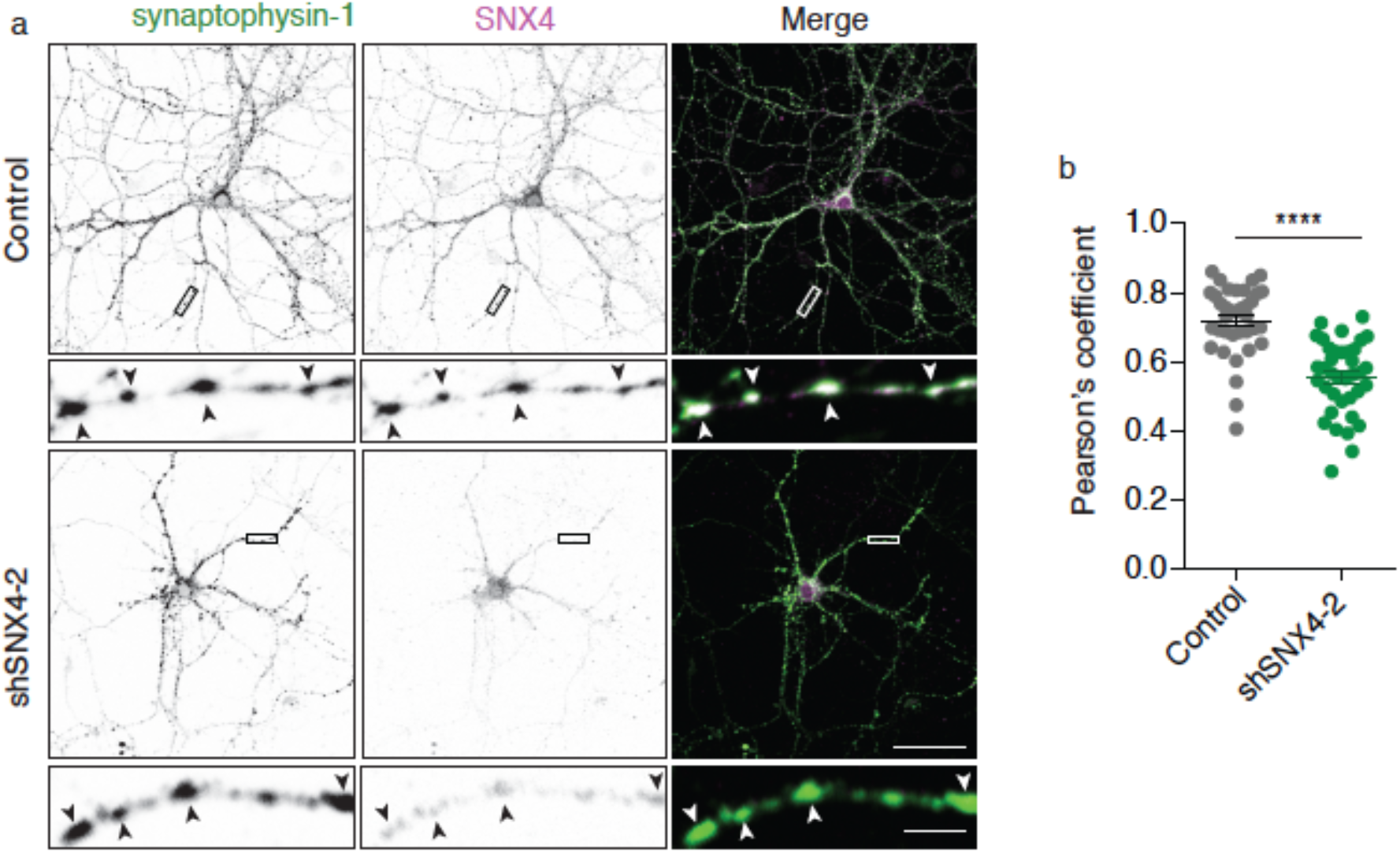
SNX4 is located to synapses. **(a)** Confocal microscopy images of hippocampal control and SNX4 knock down neurons immunolabelled with Synatophysin-1 and SNX4. Merge image of synaptophysin-1 (green) and SNX4 (magenta). Scale bar of the neuronimage=50 μm, scale bar of the zoomed neurite=5 μm. **(b)** Pearson’s coefficient for the co-localization between synatophysin-1 and SNX4 in neurites. (n=36 fields of view, N=3 animals). Detailed information (average, SEM, n and statistics) is shown in Supplementary Table S1.

To investigate the distribution of SNX4 within the synapse, we blotted subcellular fractions of the mouse hippocampus for SNX4 (Figures 4a-e). SNX4 was detected in the synaptosome (SyS) fraction, corroborating a synaptic localization of SNX4. SNX4 was also present in the synaptic membrane fraction (SyM), but was not detected in the PSD fraction (PSD). As expected, the PSD fraction was highly enriched in PSD95 and depleted for VAMP2/synaptobrevin-2. The detection of SNX4 in hippocampal subcellular fractions is thus very similar to presynaptic markers such as VAMP2/synaptobrevin-2. To verify localization of SNX4 to specific synaptic compartments, immuno-gold electron microscopy was performed using Protein A-gold 10 nm to detect the SNX4 antibody. The SNX4 immunosignal was detected inside presynaptic terminals and in the postsynaptic side, but not in the negative controls (blocking peptide, (Figure 4f’, f’’, Supplementary Figure S3). SNX4 immunosignal was more abundant in the presynaptic terminal (72.9 % of the synaptic immunogold signal) than in the postsynaptic side (27.1 % of the synaptic immunogold signal) (Figure 4k). Overall, these data show that SNX4 is present in both sides of the synapse but it is more abundant in presynaptic terminals.

**Figure 4:**
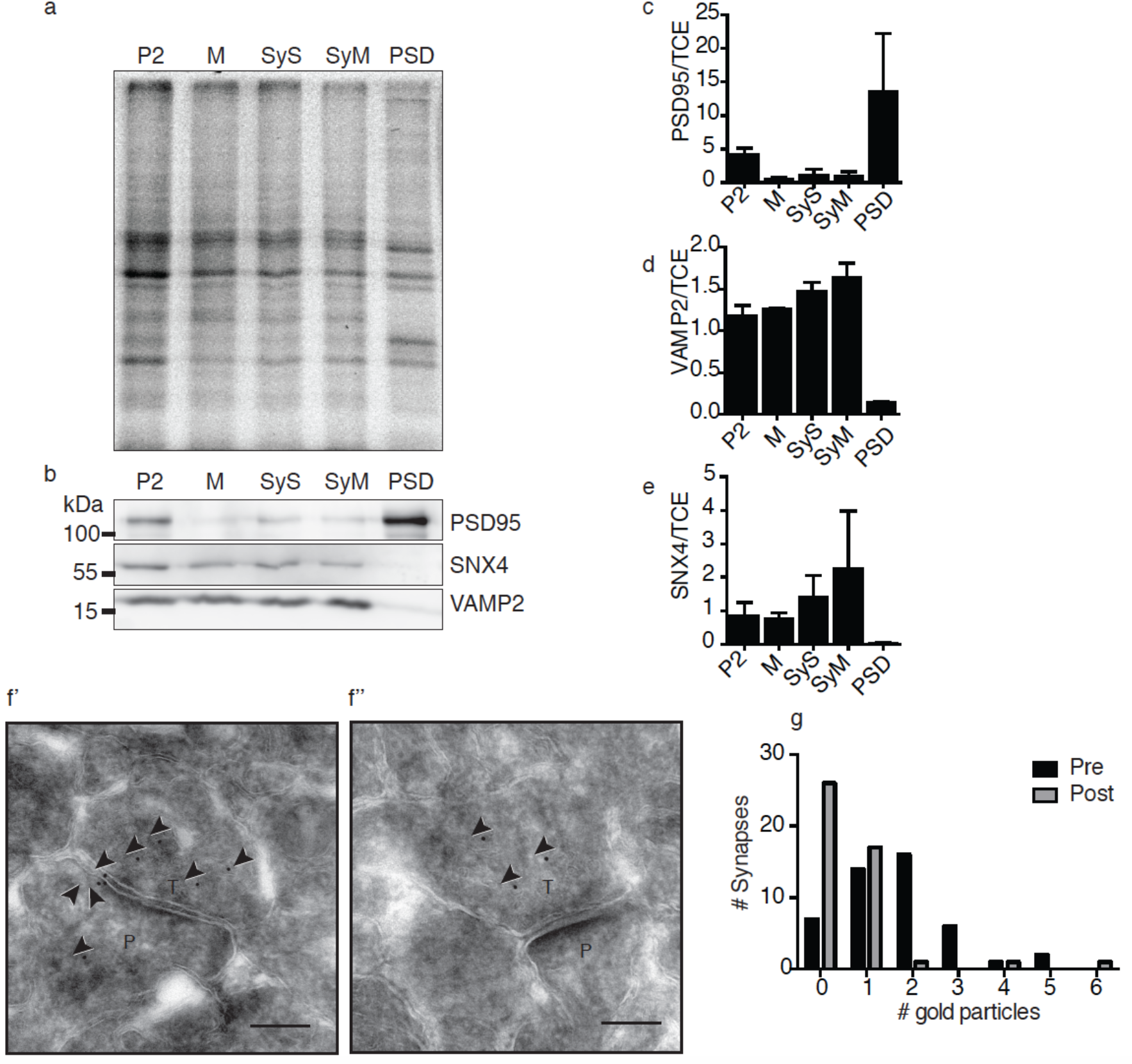
Synaptic SNX4 is predominantly located to presynaptic terminals. **(a)** Representative western blot of hippocampal subcellular fractions (pellet 2 (P2), microsomal fraction (M), synaptosomes (SyS), synaptic membrane fraction (SyM), and PSD fraction (PSD)) stained with SNX4, VAMP2/synaptobrevin-2, and PSD95. Original uncropped blots are shown in Supplementary Figure S4. **(b)** Total protein in each hippocampal subcellular fraction. Quantification of **(c)** PSD95, **(d)** VAMP2/synaptobrevin-and (e) SNX4 levels normalized to total protein. Values are presented as a ratio compared to each total hippocampus lysates. (N=3 blots/animals). (**f’,f”, f’”,f”’**) Immuno-electron micrographs of synapses labelled for SNX4 and Protein A-10nm gold conjugate. The images are representative of three independent experiments (N = 3 animals). Scale bar = 200 nm. ‘P’ indicates postsynaptic side and ‘T’ the presynaptic terminal. **(g)** Number of gold particles in the postsynaptic side and the presynaptic terminal in each synapse. (n=46 synapses, N= 3 animals). Detailed information (average, SEM, n and statistics) is shown in Supplementary Table S1.

### SNX4 depletion does not decrease TfnR in neurons, and RAB11 at synapses

In non-neuronal cells, SNX4 depletion leads to abnormal RAB11 distribution (from juxtanuclear to peripherical localization) and mis-sorting and degradation of TfnR that is recycled in these endosomes^9^. To test if this SNX4 recycling pathway is also critical for the TfnR in neurons, we measured the levels of TfnR upon SNX4 knockdown by western blot (Figure 5a, Supplementary Figure S4). Upon shSNX4 expression, TfnR levels were not changed (Figure 5b and c). We also tested if upon neuronal SNX4 depletion the distribution of RAB11 in peripheral synapses was changed. No difference was observed in the colocalization of the synaptic marker with recycling endosome marker upon SNX4 depletion, suggesting that the peripherical distribution of RAB11 is normal upon SNX4 depletion (Figure 5d and e). These data indicate that the impact of SNX4 depletion on TfnR recycling is different in neurons compared to non-neuronal cells.

**Figure 5:**
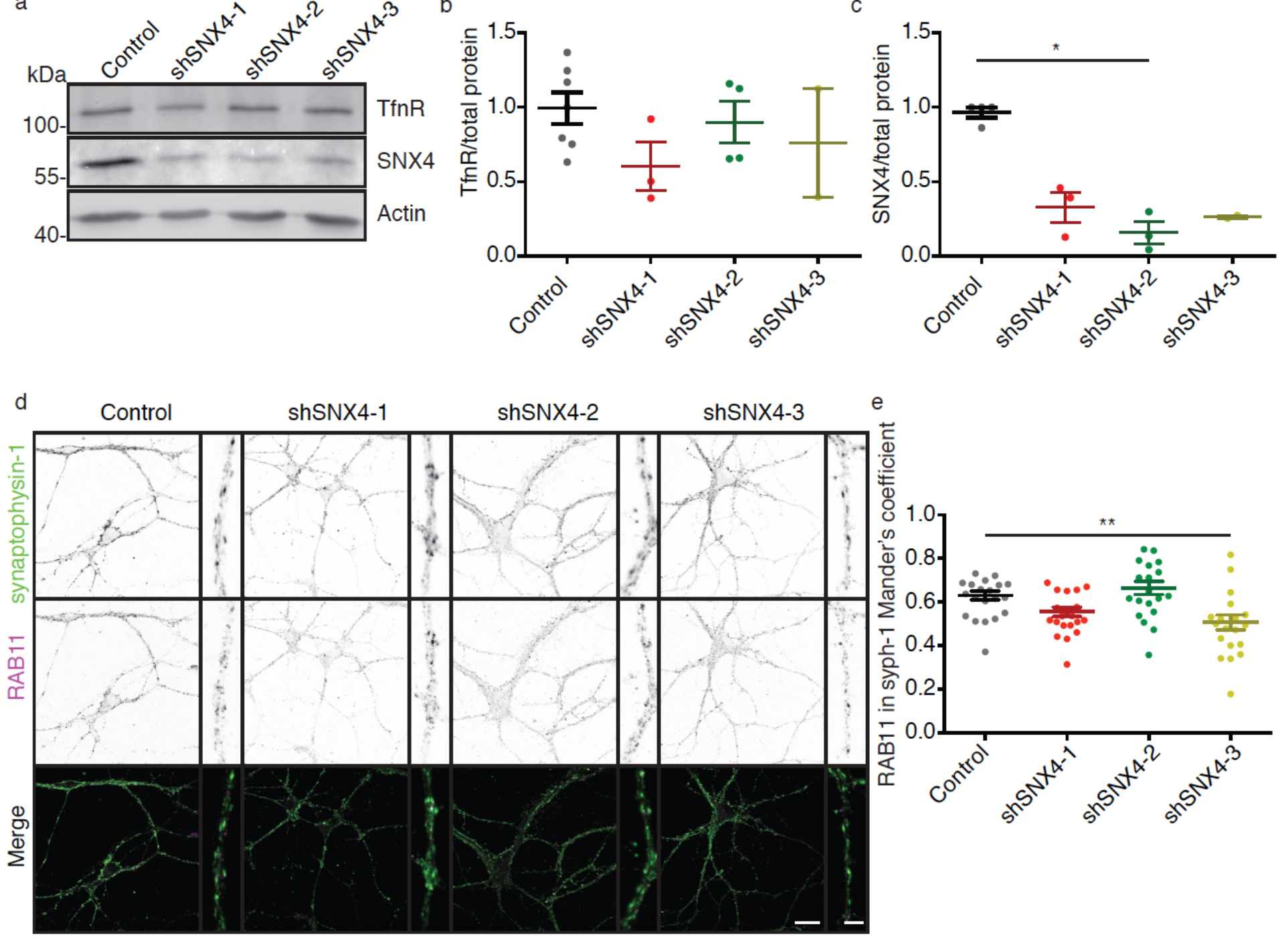
SNX4 depletion does not decrease TfnR levels in neurons and it does not decrease the recycling endosomal marker RAB11 at synapses. **(a)** Western blot of neurons infected with control shRNA (Control), and the three shRNAs against SNX4 stained for TfnR, SNX4 and actin. Original uncropped blots are shown in Supplementary Figure S3. **(b)** Quantification of TfnR levels normalized to total amount of proteins (N=3±1). **(c)** Quantification of SNX4 levels normalized to total amount of protein in western blot (N=3±1). **(d)** Confocal microscopy images of control and SNX4 KD neurons labelled with synaptophysin-1 and RAB11 antibodies. Merge image of synaptophysin-1 (green) and RAB11 (magenta). (n=21±1 neurons, N=2 animals). Scale bar of the neuron image=20 μm, scale bar of the zoomed neurite=4 μm. **(e)** Mander’s coefficient for the co-localization of synaptophysin-1 and RAB11. Detailed information (average, SEM, n and statistics) is shown in Supplementary Table S1.

### The neuronal proteome is dysregulated upon SNX4 knockdown

To explore the function of neuronal SNX4, the proteome of SNX4 knockdown and control neurons was compared. Five independent cortical cultures were infected with control shRNA and three different shRNAs against SNX4 at DIV7. At DIV15, cells were harvested and the proteins were extracted and digested into peptides for subsequent identification and quantification using LC-MS/MS^13,14^. A total of 2531 proteins were identified and quantified from 12027 peptides. Only peptides identified with high confidence were used (i.e., a Q-value ≤ 0.01 over all samples in at least one group, allowing for one outlier within each condition). Hierarchical clustering was used to classify samples into groups according to similarity. Each biological replicate from the same group (control, shSNX4-1, shSNX4-2 and shSNX4-3) clustered in separated groups (Figure 6a). Compared with control, 313, 175 and 317 proteins were uniquely dysregulated in shSNX4-1, shSNX4-2 and shSNX4-3 expressing neurons, respectively (Figure 6b). In all the three knockdown groups compared with control, 90 proteins were significantly different (Figure 6b). This suggests that the expression of each individual SNX4-targetted shRNA results in a distinct and reproducible neuronal proteome, which is difficult to explain by SNX4 knockdown alone.

**Figure 6:**
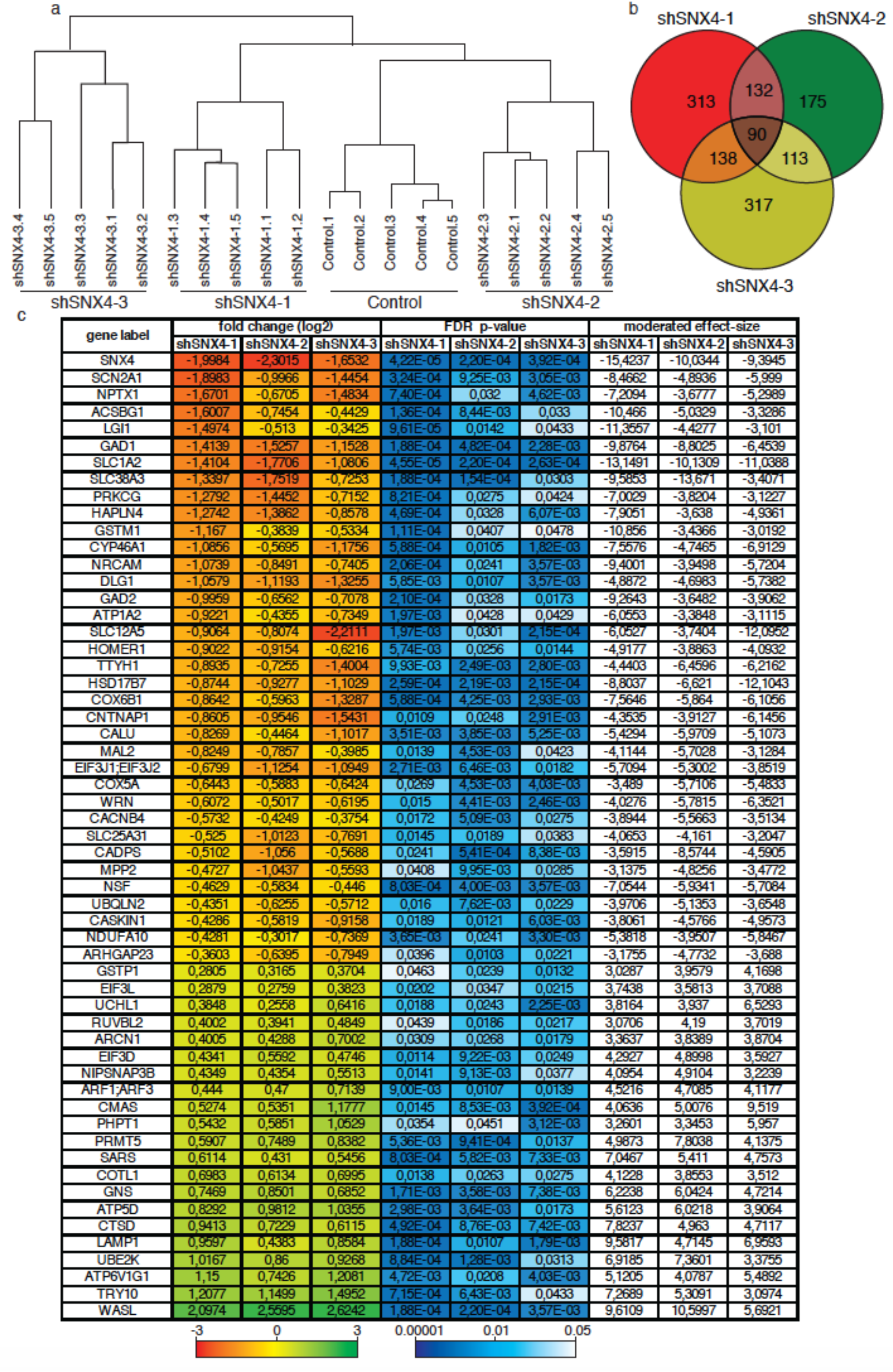
SNX4 depletion dysregulates the neuronal proteomes. **(a)** Dendrogram of the protein expression relationship between the neurons containing shRNA control and shRNA against SNX4. The hierarchical clustering reflects similarity between the samples. **(b)** Venn diagram showing the overlap among the dysregulated proteins in neurons containing shRNA against SNX4 with a moderated effect-size eBayes@limma > ±3 compared with control. **(c)** Heatmap of the protein expression of dysregulated proteins in SNX4 knock down neurons. The log2 of the fold change is color coded: Red indicates the log2 of the fold change of the maximum downregulation, green indicates the maximum upregulation and yellow no dysregulation. The p-value is color coded: Dark blue indicate the lowest p-value, and white p-value = 0.05. The moderate effect size is not color coded.

To extract functional information from the neuronal proteome upon SNX4 depletion, we focused in statistically significant proteins for which dysregulation was in same direction among the three knockdown groups (upregulated or downregulated protein in the three groups). Compared with control, 36 proteins were downregulated and 21 were upregulated in the three knockdown groups (Figure 6c). The downregulated proteins were i.e. enriched in proteins involved in *synaptic signaling* (ATP1A2, DLG1, LGI1, MPP2, SLC1A2, SLC12A5, PRKCG, GAD1, GAD2, HOMER1, CADPS, NPTX1, CACNB4), and *GABAergic synapse* functional groups (NSF, SLC12A5, PRKCG, GAD1, GAD2, SLC38A3), using g:Profiler enrichment analysis^15^.

Together, these data show that each SNX4-targetted shRNA result in a reproducible and distinct proteome, indicating potential off-target effects. Notwithstanding its limitations, these experiments show that synaptic communication-related proteins were downregulated upon three different shRNAs, suggesting a role of SNX4 in synapse-specific proteome regulation.

## DISCUSSION

A novel antibody was developed and validated for detecting endogenous mouse SNX4 by western blot, immunocytochemistry and immuno-electron microscopy. Endogenous SNX4 partially co-localized with both early and recycling endosomes in neurons, which is in accordance with the previously established role of SNX4 in non-neuronal cells^9^. Neuronal SNX4 accumulated in synaptic areas and immuno-electron microscopy revealed that SNX4 was predominantly presynaptic within the synapse. The transferrin receptor, the best described SNX4 cargo, decreases upon SNX4 depletion in nonneuronal cells^9^. In contrast, neuronal transferrin receptor levels did not change upon SNX4 depletion. Using three different shRNAs, mass spectrometry analysis revealed that synaptic communication-related proteins were downregulated upon SNX4 depletion. Hence, this study identifies neuronal SNX4 as a synaptic protein and suggests that SNX4-dependent sorting happens at synapses.

We used a shRNA approach to explore the role of SNX4 in neurons. The proteomic analysis of SNX4-depleted neurons showed that each shRNA induced a reproducible and different dysregulation of the neuronal proteome (Figure 6), while SNX4 was knocked down similarly by the three shRNAs against SNX4 (Figure 1). These uniquely dysregulated proteins are considered SNX4 depletion independent and they can be/might produce off-target effects. Off-target effects are defined as those proteins or processes which are affected by the shRNA that are independent of the target (see reviews^16,17^). Hence, in our study, only phenotypes and dysregulated proteins observed upon three independent shRNAs against SNX4 are considered SNX4 dependent. Off-target effects on neuronal morphology are common^18^, and off-target effects on neuronal migration have been also reported using nine control (scramble) shRNAs, product of an alteration of endogenous miRNA pathways^19^. Although the shRNA approach is widely used in the literature to address protein function, it needs to be tightly controlled to draw solid conclusions. Functional assays require a rescue experiment using shRNA-resistant constructs to assure the effect depends on the target.

Most SNX4 studies have been performed in mitotic cells, where SNX4 localizes in early and recycling endosomes^5,7,9,20,21^. In neurons, we found that SNX4 also co-localized with early (RAB5) and recycling (RAB11) endosomal markers, suggesting that neuronal SNX4 functions in the previously described recycling pathway from early endosome to the plasma membrane through the recycling endosome. In HeLa cells, SNX4 silencing leads to abnormal RAB11 puncta distribution: from juxtanuclear to peripherical distribution^9^. In neurons, SNX4 depletion did not affect the RAB11 levels (Figure 2d, e) nor the distribution of RAB11 puncta in synapses, indicating that SNX4 depletion does not affect the recycling endosome distribution to synapses (Figure 5). The best described SNX4-dependent recycling cargo is the TfnR, which is decreased upon SNX4 depletion in HeLa cells^9^. In neurons, SNX4 depletion did not decrease TfnR levels measured by western blot and confirmed by mass spectrometry (Figure 5). The impact of SNX4 depletion on TfnR levels in neurons is different compared with HeLa cells, but it is not possible to exclude that the TfnR is not recycled by SNX4. For example, neuronal SNX4-dependent cargo might not be degraded in lysosomes upon SNX4 depletion, but be mis-trafficked or accumulated in internal compartments where it cannot function without affecting the total protein level. Nevertheless, SNX4-dependent recycling seems to be different in different neurons, which may reflect a different demand of these large, polarized, and post-mitotic cells specialize in neurotransmission.

In HeLa cells, SNX4 recycles cargo proteins from the early endosome back to the plasma membrane through recycling endosomes, avoiding its lysosomal degradation. Hence, when SNX4 is depleted, SNX4-dependent cargo is degraded at the lysosome^9^. Upon SNX4 depletion, we found that the expression of some proteins involved in synaptic transmission were decreased. Although lysosomal processes were not detected as an affected functional protein group, the individual lysosomal proteins Cathepsin D and LAMP1 were increased upon SNX4 knockdown, which might suggest an increase lysosomal degradation demand (Figure 6). Potentially, the SNX4 recycling pathway is also present in synaptic terminals where it recycles synaptic proteins and regulates the local synaptic proteome.

In neurons, SNX4 has been studied in the context of Alzheimer’s disease. SNX4 depletion decreased APP^10^ levels, increased BACE1 in late endosomal compartments^10,12^, and modulated Aβ production^12^. In our mass spectrometry data set, BACE1 was not detected and APP was identified using only one peptide which overlaps with the sequences of Aβ, thus, these two cannot be distinguished. APP levels were decreased upon shSNX4-3 expression, but not upon shSNX4-1 and shSNX4-2 (shSNX4-3 p=0.0172), highlighting again the importance of independent shRNAs.

Endogenous SNX4 was expressed in all tested brain areas (Supplementary Figure 2), indicating that SNX4 function is important for all brain regions, in line with its evolutionary conservation across eukaryotic organisms^1,2^. In neurons, endogenous SNX4 highly colocalized with synaptic markers (Figure 3 and Supplementary Table S1) and was found in synaptic fractions. Although the new antibody showed an unspecific band in western blot analyses, immunocytochemistry-labeling of SNX4 in synapses proved to be specific: the signal was decreased upon SNX4 shRNAs and restored upon introduction of a shRNA-resistant SNX4 variants. Immuno-electron microscopy revealed that endogenous SNX4 is present in both the post- and pre-synaptic terminals. Hence, SNX4 is a novel synaptic protein, which suggests that SNX4-mediaded recycling may be required for synaptic function. Although, presynaptic SNX4 unlikely traffics the TfnR, synaptic SNX4 might be involved in trafficking proteins to the synaptic plasma membrane. In presynaptic terminals, the endosomal system is known to be involved in the insertion of metabotropic receptors such as G protein–coupled receptors (GPCRs) (see review^23^). GPCRs can be localized at presynaptic terminals and are important regulators of synaptic communication (see review^24^). On the postsynaptic side, the endosomal system is also involved in the insertion of neurotransmitter receptors in the plasma membrane, which is an also known mechanism of synaptic plasticity (see review^25^). Hence, SNX4 might play a role in synaptic endosomal sorting which may regulate synaptic plasticity.

Although endosomal sorting genes are ubiquitously expressed, mutations in these genes are notably associated with neurodegenerative diseases^26^. A hallmark that occurs early in neurogenerative diseases progression is synapse loss^27^. Therefore, the novel identification of SNX4 at synaptic terminals opens a new line of research on the role of this protein at synapses. Addressing synaptic endosomal sorting might provide a better understanding of the pathogenesis of neurodegenerative disorders, thereby providing potential targets for therapeutic intervention at an early stage of disease progression that precedes irreversible cell death.

## MATERIALS AND METHODS

### Plasmids

Short harping RNA (shRNA) were cloned in to a lentiviral expression vector under the U6 promotor. To report lentiviral infection the plasmid also contained mCherry under synapsin promotor. The target sequences of the shRNAs were as follows: GGG AAT GAC TAC CAA ACT C (shSNX4-1), GCA GTG GAA TAG ATA CAT TAT (shSNX4-2), GCT GAT ATT GAA CGC TTC AAA (shSNX4-3), TTC TCC GAA CGT GTC ACG T (shControl, scramble)^28^. Mouse SNX4 cDNA was used to induce the silence mutagenesis in rescue constructs. The following forward primers were used: GAAGGGAATGACAACGAAGCTTTTTGGTCAAGAAACTCCAG (shSNX4-1) GGGCTGATATCGAGCGCTTTAAAGAACAAAAG (shSNX4-3).

### Laboratory animals

Animal experiments were approved by the animal ethical committee of the VU University/VU University Medical Centre (“Dier ethische commissie (DEC)”; license number: FGA 11-03) and, they are in according to institutional and Dutch governmental guidelines and regulations.

### Primary cell culture

Primary neurons were cultured from mouse E18 hippocampi or cortices. Briefly, tissue was dissected in Hanks balance salt solution (HBSS, Sigma) with 10mM HEPES (Life Technologies) and digested by 0.25% trypsin (20 minutes at 37°C; Life technologies) in HBSS. The tissue disassociation was performed with fire-polished Pasteur pipettes in DMEM with FCS. The neurons were spun down and re-suspended in neurobasal medium with 2% B-27, 18 mM HEPES, 0.25% glutamax and 0.1% Pen-Strep (Life Technologies). Neurons were plated in coated coverslips with poly-L-ornithine (PLO, Sigma) and laminin (Sigma), or astrocyte monolayer. Neurons were maintained at 37°C and 5% CO2 until the day of the experiment.

### Subcellular fractioning

Subcellular fractions were obtained from hippocampi from three-month-old C57BL6 mice as previously described^29,30^. Isolated hippocampi were homogenized on a dounce homogenizer (potterS; 12 strokes, 900 rpm) using homogenizer buffer (0.32 M Sucrose, 5 mM HEPES pH 7.4, Protease inhibitor cocktail (Roche)), and spun at 1000xg for 10 minutes at 4°C to obtain Supernatant 1 (S1). S1 was centrifuged at 20,000xg for 20 minutes to obtain pellet 2 (P2) and supernatant 2 (S2). S2 was ultracentrifuged at 100,000xg for 2 hours to obtain the pellet containing the microsomal fraction (M). S1 was ultracentrifugated in a 0.85/1.2 M sucrose density gradient at 100,000xg for 2 hours to obtain Synaptosomes (SyS) at the interface of 0.85/1.2M sucrose. SyS were exposed to a hypotonic shock of 5 mM HEPES pH 7.4 with protease inhibitor for 15 minutes, and sucrose gradient ultracentrifugated as stated above to obtain the synaptic membrane fraction (SyM) at the interface of 0.85/1.2M. SyS was also treated with 1% Tx-100 for 30 minutes, layered on top of 1.2/1.5/2M sucrose, centrifuged at 100,000xg for 2 hours, to obtain the PSD fraction (PSD) at the interface of 1.5/2M sucrose.

### Western Blot

Wild type mouse brain regions were homogenized in ice-cold PBS with protease inhibitors and lysed (100μl Laemmli sample buffer (2% w/v sodium dodecyl sulfate (SDS), 10 % v/v Glycerol, 0.26 M β-mercaptoethanol, 60 mM Tris-HCl pH 6.8, and 0.01% w/v Bromophenolblue) per each mg). DIV14-15 cortical neurons were washed with ice-cold phosphate-buffered saline (PBS), scraped, lysed in Laemmli sample buffer. Samples were boiled for 10 minutes at 90°C, loaded in SDS-PAGE (10% 1 mm acrylamide gel with 2,2,2-Trichloroethanol) and, transferred into Polyvinylideenfluoride (PVDF) membranes (Bio-rad) (1 hour, 0.3 mA, 4°C). Membranes were blocked using 2% milk (Merck) with 0.05% of normal goat serum (NGS) in PBS-T (PBS with 0.1% Tween-20), incubated overnight at 4°C with the primary antibodies in PBS-T (Supplementary Table S2), with secondary alkaline phosphatase conjugated antibodies (1:10000, Jackson ImmunoResearch) in PBS-T during 1 hour at 4°C, and incubated 5 minutes with AttoPhos (Promega). Images were acquired with a FLA-5000 fluorescent image analyzer (Fujifilm) and analyzed with ImageJ Gel Analysis.

### Immunocytochemistry and Confocal Imaging

Neurons at DIV 14-15 were fixed with 2% paraformaldehyde in PBS and cell culture media for 10 minutes followed by 4% paraformaldehyde in PBS for 30 minutes at room temperature. Then, neurons were washed, permeabilized with 0.5% Triton X-100 for 5 minutes, blocked with 2% normal goat serum and 0.1% Triton X-100 in PBS for 40 minutes, incubated 1 hour with primary antibodies (Supplementary Table S2), 1 hour with secondary antibodies conjugated to Alexa dyes (1:1000, Molecular Probes), and mounted on microscope slides with Dabco-Mowiol at room temperature. Images were acquired using a Carl Zeiss LSM510 meta confocal microscope with a Plan-Neofluar 40x/1.3 oil objective and a Nikon Eclipse Ti with 63x/1.4 oil objective controlled by NisElements 4.30 software. Colocalization was analyzed using JACoB plugin^31^. For quantification of protein levels, intensity was measure inside a neuronal mask in ImageJ.

### Electron microscopy

Hippocampi of 2 months old mice were fixed in 4% PFA with 0.1% glutaraldehyde in 0.1M PB and embedded in increasing concentrations of gelatin at 37°C (5 minutes 2% gelatin, 15 minutes 5% gelatin, 30 minutes 10% gelatin, 10 minutes 12% gelatin, 60 minutes 12% gelatin). The hippocampi were infiltrated in 2.3 M sucrose at 4°C and frozen in liquid nitrogen. Seventy nm thick sections were obtained with a cryo-ultramicrotome (UC6, Leica), collected at −120°C in 1% methyl-cellulose and 1.2 M sucrose and transferred onto formvar/carbon-coated copper mesh grids. The sections were washed with PBS at 37°C, treated with 0.1% glycine, blocked with 0.1% of BSA and 0.1% cold water fish gelatin, incubated during 2 hours with SNX4 antibody in blocking solution (Supplementary Table S2), and 1 hour with Protein A-10 nm gold (1: 25, CMC, UMC Utrecht, Netherlands) at room temperature. The sections were counterstained with 0.4% uranyl acetate in 1.8% methyl-cellulose on ice and imaged on a Tecnai 12 Biotwin transmission electron microscope (FEI company).

### Proteomics

DIV14-15 cortical neurons (250.000 neurons/mL in laminin/poly-L-ornithine coated 6-well plates) were washed and scraped with ice cold PBS with protease inhibitor cocktail (Roche). Neurons were pelleted (5 minutes at 3000 g at 4°C), lysed in loading buffer (0.05 M Tris-HCl pH 6.8, 2% SDS, 10% glycerol, 0.1M DTT, 0.001% bromophenol), boiled at 90°C for 5min, and incubated with acrylamide at room temperature for 30 minutes. Each sample (~500.000 neurons normalized to the total amount of proteins) was separated about 1cm on a 10% SDS polyacrylamide gel, fixed overnight and stained with colloidal Coomassie Brilliant Blue G. Each sample lane was cut into small fragments and transferred to the wells of a MultiScreen-HV 96 well filter-plate. Destined and dried fragments were re-swelled with Trypsin/Lys-C Mix solution (Promega) overnight in a humidified chamber at 37°C. The peptides were extracted from the gel pieces with 50% acetonitrile in 0.1% TFA, and then with 80% acetonitrile in 0.1% TFA. Finally, the peptides were dried in solution using a speedvac and stored at −20°C.

The peptides were re-dissolved in 2% acetonitrile/0.1% formic acid solution containing iRT reference peptides and injected into the Ultimate 3000 LC system. The peptides were trapped on a 5 mm C18 PepMap 100 column for 5 minutes and separated on a homemade 200 mm C18 Alltima column. The reverse phase liquid chromatography was performed by linearly increasing the acetonitrile concentration in the mobile phase at a flow rate of 5 μL/minute: from 5 to 22% in 88 minutes, to 25% at 98 minutes, to 40% at 108 minutes and to 95% in 2 minutes. The separated peptides were electro-sprayed into the TripleTOF 5600 MS (Sciex) with a micro-spray needle (at a voltage of 5500 V). The mass spectrometer was set in data-independent acquisition at high sensitivity and positive mode under the following parameters: parent ion scan of 100 msec (mass range of 350-1250 Da), SWATH mass range between 450-770 m/z, SWATH window of 8 Da, MS/MS scan time of 80 msec per window (range 200-1800 Da), collision energy for each window was determine for a 2+ ion centered upon the window, with a spread of 15 eV.

Data was analyzed using Spectronaut 8.0^32^ and a spectral library created from merging two data-dependent analyses of wild type hippocampal neuron cultures and hippocampal synaptosomes containing spike-in iRT peptides from Biognosys^33^. The retention time prediction was set to dynamic iRT; the cross-run normalization based on total peak areas was enabled. The resulted peptide abundances were processed using R language for statistical computation. Protein abundances were computed using Spectronaut normalized peak area, and Loess normalized using the ‘normalizeCyclicLoess’ function from limma R package (fast method and 10 iterations)^34^. Empirical Bayes moderated t-statistics with multiple testing correction by false discovery rate (FDR) was performed on log-transformed protein abundances as implemented by the ‘eBayes’ and ‘topTable’ functions from limma R package. Only proteins fulfilling the following criteria were used for further analysis: moderated effect-size eBayes@limma was superior to ±3, direction of the dysregulation was equal among the SNX4 knockdown groups, and Empirical Bayes moderated t-statistics FDR was ≤ 0.05. These proteins were analyzed in g:Profiler (version: r1741_e90_eg37) using the total identified proteins with high confidence as a gen list background, and using default setting (including *Homo sapiens* as a default organism)^15^.

### Statistical Analysis

Data are expressed as mean values ± standard error of the mean (SEM). The Shapiro-Wilk normality test was used to evaluate the distribution of the data. Bartlett’s test was used to test homoscedasticity. If comparing two homoscedastic and normal distributed groups, t-test was used, otherwise, Mann-Whitney test was used. If comparing more than two groups, data were normally distributed and homoscedastic, data were compared by one-way analysis of variance (ANOVA). Dunnets post-hoc tests were performed after a significant effect was detected by comparing the different knockdown groups to the control. In case of comparing more than two groups, data were not normality distributed and homoscedastic, the Kruskal-Wallis test was used with Dunn’s multiple test as post-hoc. When P-values were lower than 0.05, significance was noted in the figure as: *P<0.05, **P<0.01, ***P<0.001, ****P<0.0001.

## Data availability

The datasets generated and analyzed during the current study are available from the corresponding author on request.

## AUTHOR CONTRIBUTIONS

S.V.S. performed experiments and analyzed the data. A.W. collected and analyzed confocal images for the SNX4 colocalization studies. M.A.G.L. and K.W.L produced and critically discussed the proteomic data. S.V.S. and J.R.T.vW. designed the experiments and, wrote the manuscript.

## ACKNOWLEDGMENTS

The authors thank Prof. Dr. Matthijs Verhage and Prof. Dr. Peter J. Cullen for their critical reading of the manuscript, Joke Wortel for housing and breeding the mice, Frank den Oudsten and Desiree Schut for providing cell cultures, and Robbert Zalm and Joost Hoetjes for cloning and lentiviral production, Rozemarijn Jongeneel for confocal analysis and Joke Wortel and Marien P. Dekker for assistance with EM analysis. EM analysis was performed at the VU/VUmc EM facility (ZonMW 91111009). This work was supported by the EC under FP7-PEOPLE-2013 (607508) and Alzheimer Nederland (WE.03_2016-05).

## SUPPLEMENTARY FIGURES LEGENDS

**Supplementary Figure S1:**
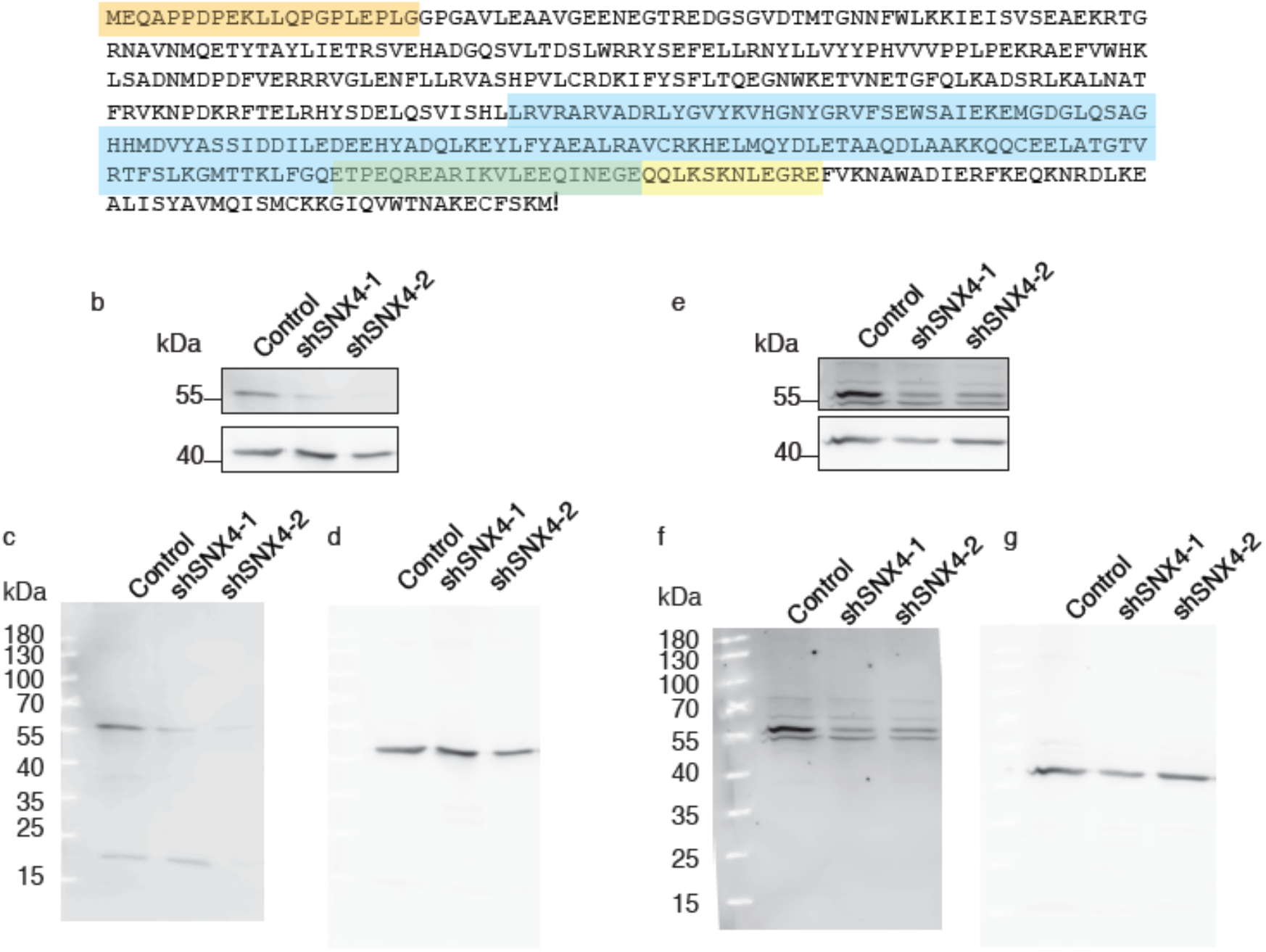
Epitopes of the different antibodies against SNX4. **(a)** Sequence of amino acids of mouse SNX4 (>gi|18017596|ref|NP_542124.1| sorting nexin-4 *[Mus musculus]).* The epitopes of the different antibodies are highlighted. In orange, the epitope of SNX4 antibody from cat. N. 392 003, Synaptic Systems (1-21 amino acids of mouse SNX4). In blue, epitope from cat. N. HPA005709, Sigma (238-386 amino acids of human SNX4). In yellow, epitope from cat. N. sc-271403, Santa Cruz (361-393 amino acids of human SNX4). The green is just the product of the overlapping sequences highlighted in yellow and blue. **(b)** Representative western blot of control neurons and neurons with shRNAs against SNX4 stained for SNX4 (N. sc-271403, Santa Cruz) and actin. Original uncropped blots for SNX4 (N. sc-271403, Santa Cruz) **(c)** and acting **(d). (e)** Representative western blot of control neurons and neurons with shRNAs against SNX4 stained for SNX4 (N. HPA005709, Sigma) and actin. Original uncropped blots for SNX4 (N. HPA005709, S) **(f)** and actin **(g)**.

**Supplementary Figure S2:**
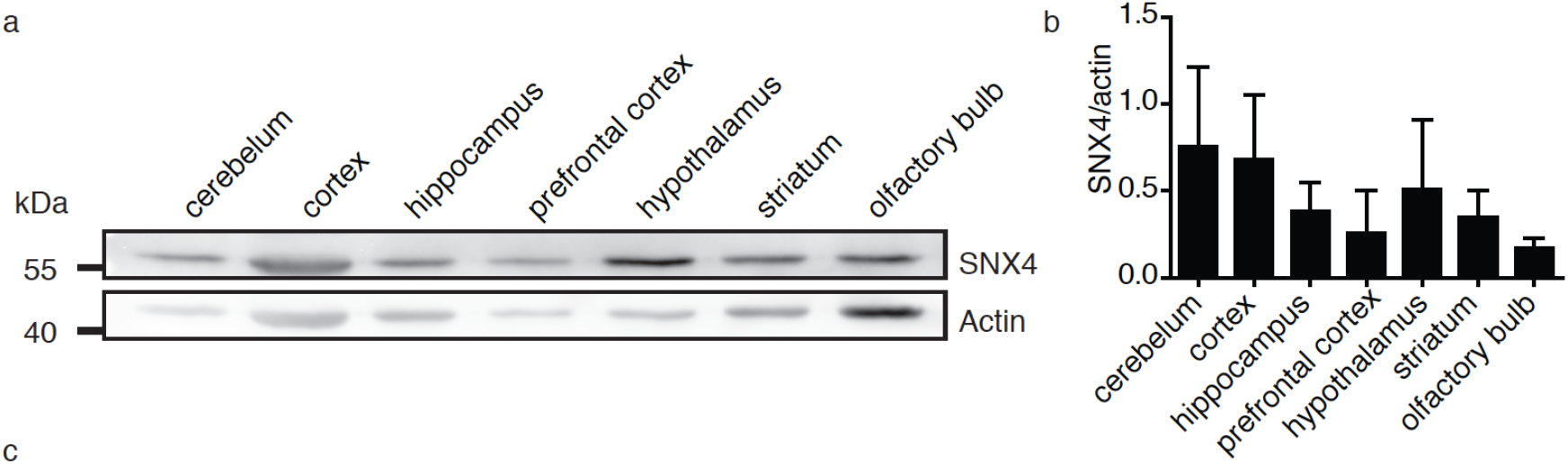
SNX4 is expressed in different brain regions. **(a)** Western blot of different mouse brain areas for SNX4 and actin. Original uncropped blots are shown in Supplementary Figure S4. **(b)**. Quantification of SNX4 levels normalized to actin in western blot. (N=2 or 3 blots/animals). Detailed information (average, SEM, and n/N) is shown in Supplementary Table S1.

**Supplementary Figure S3:**
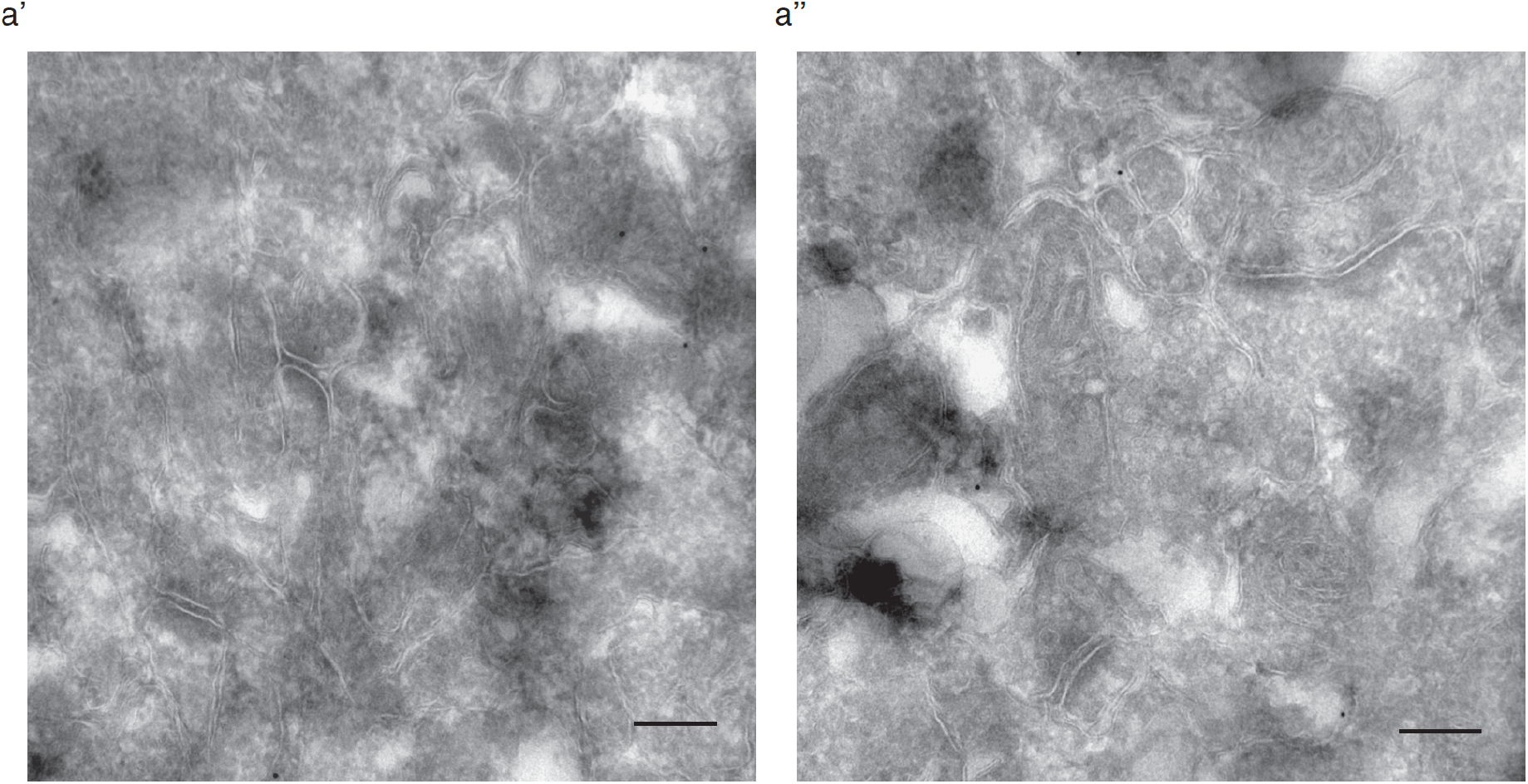
Electron micrographs of the negative controls for immuno-gold labelling against SNX4 (a’, a’’). Negative control processed in parallel with the immunolabelling with SNX4 but preincubating the primary antibody with the blocking peptide (Synaptic Systems, Cat. No. 392-0P at a ratio of 1:10). Scale bar=200nm.

**Supplementary Figure S4:**
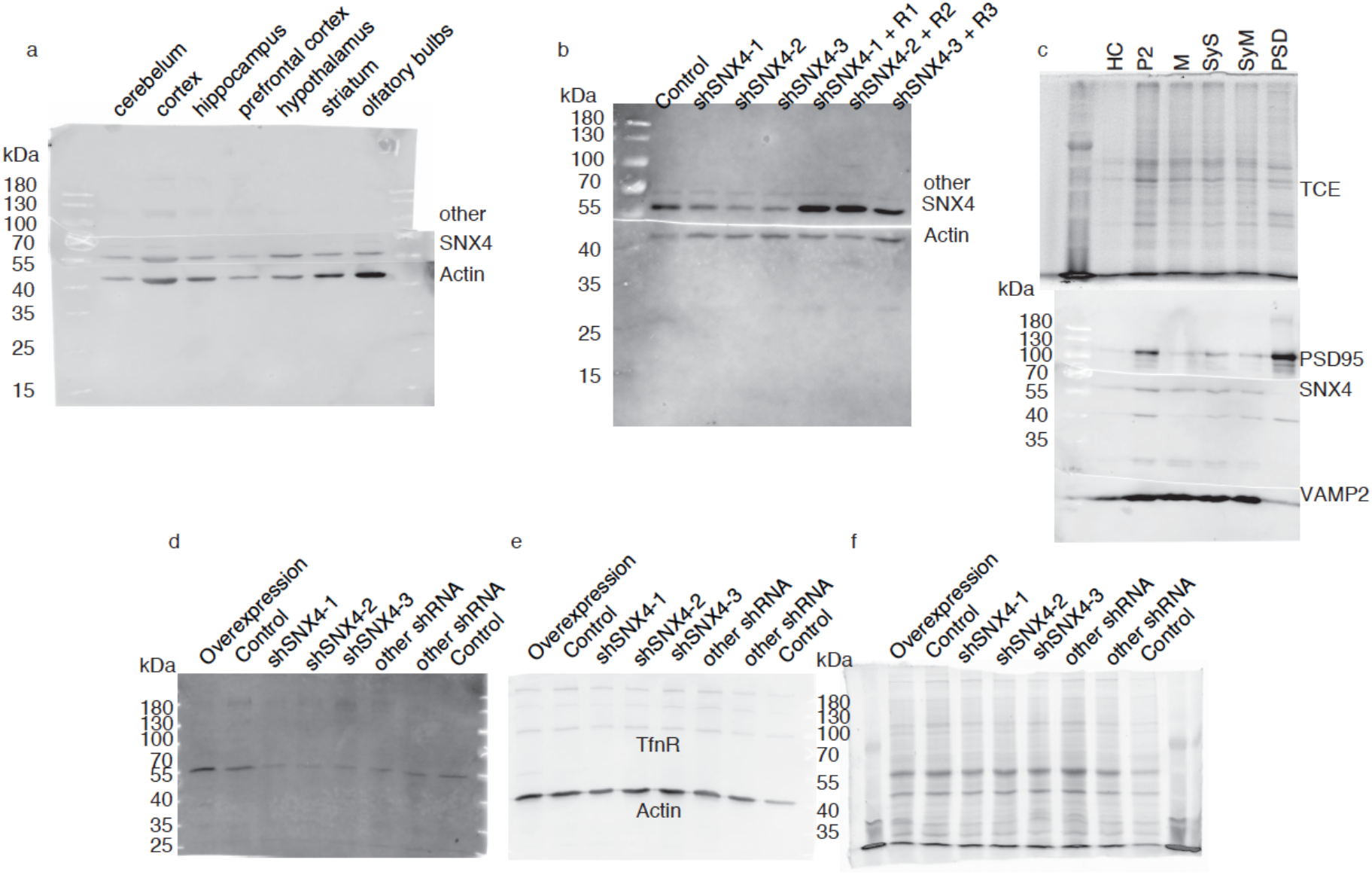
Original uncropped blots. **(a)** Original uncropped blots for SNX4 and actin of the data shown in Supplementary Figure S2. The top part was blotted for Fbxo41 which is not relevant for this study. **(b)** Reblot for actin (low part of the blot) and for a non-relevant antibody (upper part of the blot) from Figure 2c. **(c)** Gel stained with TCE of the data shown in Figure 4 and original uncropped blots for PSD95, SNX4, and VAMP2/synaptobrevin-2. The first line is full hippocampal lysate from which the subcellular fractions were obtained. Original uncropped western blot of the data shown in Figure 5a stained for SNX4 **(d)** and for **(e)** TfnR and actin and **(f)** gel stained with TCE.

**Supplementary Figure S5:**
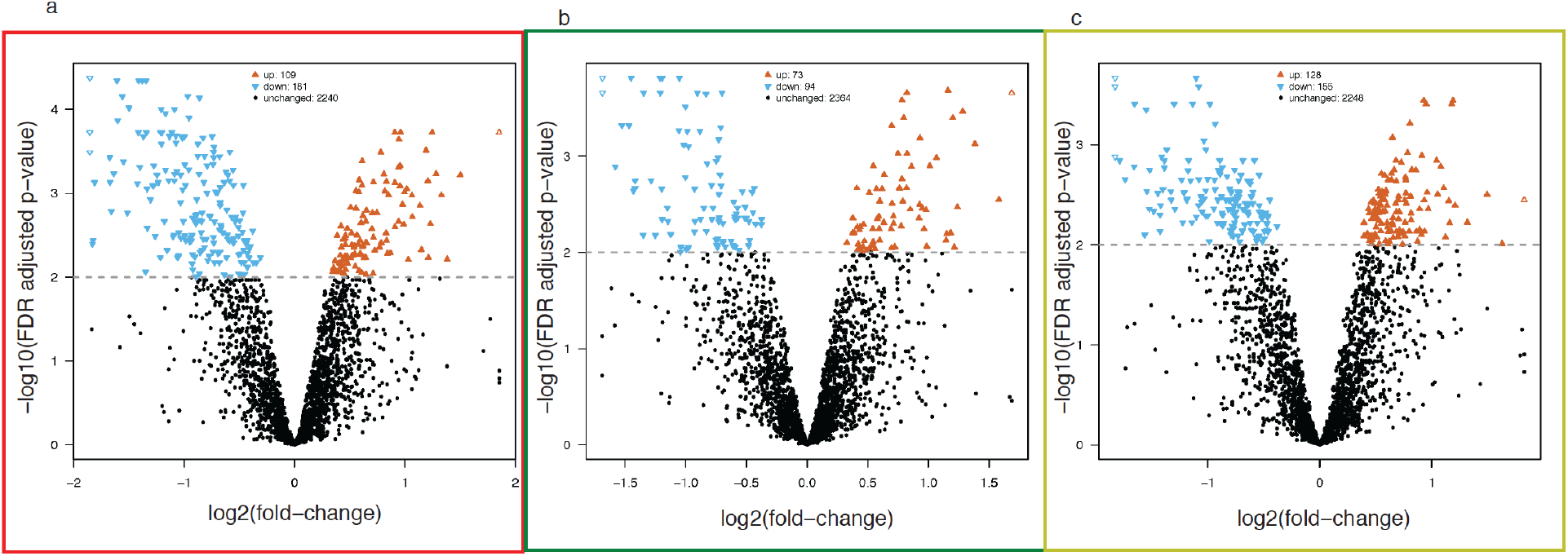
Volcano plots showing the distribution of protein expression in cortical neurons expressing shRNA against SNX4 compared with control neurons. shSNX4-1 **(a)**, shSNX4-2 **(b)**, and shSNX4-3 **(c)**.

**Supplementary Figure S6:**
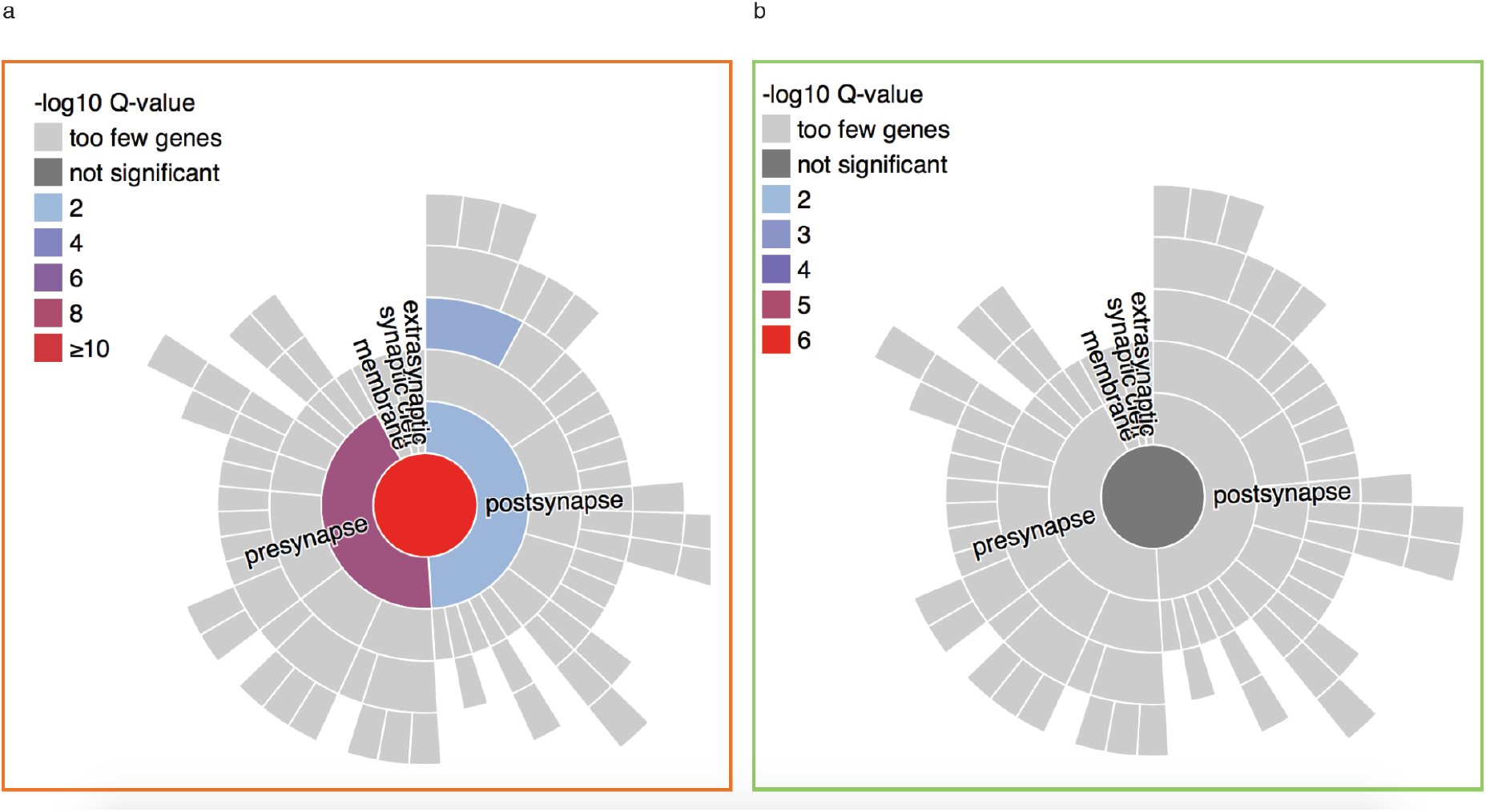
Enrichment analysis upon shRNA against SNX4 expression by SynGO. Downregulated **(a)**, and upregulated **(b)**.

## Supplementary Tables

**Supplementary Table S1: Summary of the mean, SEM, n/N numbers and statistics of measured variables in the study**. Not applicable (empty cells).

**Supplementary Table S2: Primary antibodies specifications and concentrations.**

